# Robustness of trinucleosome compaction to A-tract mediated linker histone orientation

**DOI:** 10.1101/2021.08.13.456082

**Authors:** Madhura De, Martin Würtz, Gabriele Müller, Katalin Tóth, Rebecca C. Wade

**Affiliations:** Division of Biophysics of Macromolecules, German Cancer Research Center (DKFZ), 69120 Heidelberg, Germany; Molecular and Cellular Modelling Group, Heidelberg Institute for Theoretical Studies (HITS), 69118 Heidelberg, Germany; Faculty of Biosciences, Heidelberg University, 69120 Heidelberg, Germany; Center for Molecular Biology (ZMBH), DKFZ-ZMBH Alliance, Heidelberg University, 69120 Heidelberg, Germany; Department of Biophysics and Cell Biology, Faculty of Medicine, University of Debrecen, Hungary; Interdisciplinary Center for Scientific Computing (IWR), Heidelberg University, 69120 Heidelberg, Germany; Department of Biological Chemistry and Molecular Pharmacology, Harvard Medical School, Boston, MA 02115, USA

**Author notes:** To whom correspondence should be addressed. Katalin Tóth. To whom correspondence should be addressed. Rebecca C. Wade; @Rebecca_Wade_C.

**Keywords:** chromatin structure, linker histone, nucleosome, single-pair FRET, A-tract DNA

## Abstract

Linker histones (LH) have been shown to preferentially bind to AT-rich DNA, particularly A-tracts, contiguous stretches of adenines. Using spFRET (single pair Förster/Fluorescence Resonance Energy Transfer), we recently found that the globular domain (gH) of *Xenopus laevis* H1.0b LH orients towards A-tracts on the linker-DNA (L-DNA) while binding on-dyad in LH:mononucleosome complexes. Here, we investigate the impact of this A-tract-mediated orientation of the gH on the compaction of higher-order structures by studying trinucleosomes as minimal models for chromatin. Two 600 bp DNA sequences were constructed, each containing three consecutive Widom 601 core sequences connected by about 40 bp L-DNA but differing in the positioning of A-tracts on either the outer or the inner L-DNAs flanking the first and third Widom 601 sequences. The two inner L-DNAs were fluorescently labelled at their midpoints. Trinucleosomes were reconstituted using the doubly labelled DNA, core histone octamers and H1.0b. SpFRET was performed for a range of NaCl concentrations to measure the compaction and whether gH orientations affected the stability of the trinucleosomes to salt-induced dissociation. While the LH compacted the trinucleosomes, the extent of compaction and the stability were similar for the two DNA sequences. Modeling constrained by the measured FRET efficiency suggests that the structures adopted by the trinucleosomes correspond to the standard zig-zagged two-helical start arrangement with the first and third nucleosomes stacked on top of each other. In this arrangement, the first and third LHs are insufficiently close to interact and affect compaction. Thus, despite differences in the positioning of the A-tracts in the sequences studied, LH binding compacts the corresponding trinucleosomes similarly.

**Why it matters:** The compaction and three-dimensional structure of chromatin affect the exposure of the DNA and thus regulate gene expression. Linker histone proteins bind to nucleosomes and thereby contribute to chromatin compaction. We here investigated whether the DNA A-tract-mediated orientation of a linker histone globular domain affects chromatin structure by using a trinucleosome as a minimal model for chromatin. Our observations suggest that the trinucleosome structure and compaction are robust against differences in linker histone globular domain orientations.

**eTOC blurb:** We investigate whether DNA sequences, such as adenine-tracts, and sequence-induced linker histone reorientation affect chromatin structure. Using trinucleosomes as model systems for chromatin, we demonstrate that the chromatin structure and compaction are robust to the studied DNA sequence differences and sequence-induced linker histone orientation.

## Introduction

The binding of the highly positively charged linker histone (LH) H1 protein to the nucleosome plays an important role in the compaction of chromatin (1). The LH associates with the nucleosome in the region bounded by the linker-DNA (L-DNA) arms entering and exiting the nucleosome core particle (2–6). The LH has a tripartite structure composed of a conserved globular domain (gH), a 100 residue long, intrinsically disordered C-terminal domain (CTD) and a shorter N-terminal domain (7). The CTD has been observed experimentally to associate with one (5, 8) or both L-DNA arms (6), and in a salt-dependent manner with one (5 and 15 mM NaCl) or both (80 and 150 mM NaCl) L-DNAs in coarse-grained simulations (9). The association of the CTD with the L-DNAs has also been suggested to have an effect on the charge distributions of nucleosomal arrays (5). The gH has been observed to bind the nucleosome (10, 11) in an on-dyad (3–6) or an off-dyad (8, 12) position, or a dynamic combination of the two (13). The positioning of the gH has been suggested, from both experimental (7,9) and computational studies (15), to affect the compaction of nucleosomal arrays. In this work, we investigate whether the orientation in which the gH binds on-dyad to the nucleosome affects chromatin compaction.

To address this question, the gH of LHs should be made to associate with the nucleosomes in a specific manner, either by mutating the gH domain residues that determine on- or off-dyad binding (12), or by exploiting the preference of the LH gH for AT-rich DNA or A-tracts (a series of contiguous adenines) (6, 16–27). Here, we built on our earlier study of mononucleosome-LH complexes by spFRET (27) that showed that the gH domain of a full-length LH recognizes an 11 bp A-tract on the L-DNA by binding on-dyad in a specific orientation (Figure 1B ii). We therefore here addressed the question of whether such an A-tract mediated orientation of the gH domain affects the compaction of nucleosomal arrays. For this purpose, we constructed two trinucleosomes. Each trinucleosome had a 600 bp DNA sequence with three consecutive Widom 601 (strongly nucleosome positioning) sequences (28) situated about 40 bp apart from each other, thereby resulting in a nucleosome repeat length (NRL) of 187 bp. The flank sequences of nucleosomes (Nuc) 1 and 3 were either A-tract (11 contiguous adenines complementary to thymines) or purely GC-tract. Depending upon the location of the A-tracts (Table 1), the trinucleosome constructs were named A-far (A-tracts on outer L-DNAs and GC-tracts on inner L-DNAs) or A-near (A-tracts on inner L-DNAs and GC-tracts on outer L-DNAs). Both the constructs were doubly labelled at the mid-points of the two inner L-DNAs with donor (Alexa 488) and acceptor (Alexa 594) fluorophores (Figure 1A). By performing single-pair FRET spectroscopy and structural modeling, we were able to investigate whether or not the positioning of A-tracts in nucleosomal arrays affects the extent of LH-dependent compaction of these arrays.

**Table 1:** Flank sequences of the outer and inner L-DNA arms for the A-near and A-far trinucleosomes.

| Trinucleosome construct | Inner L-DNA flank sequence | Outer L-DNA flank sequence |
| --- | --- | --- |
| A-near | 5' - AAAAAAAAAA -3' | 5' - GGGCGGCCGCG -3' |
|  | 3' - TTTTTTTTTT -5' | 3' - CCCGCCGGCGC -5' |
| A-far | 5' - GGGCGGCCGCG -3' | 5' - AAAAAAAAAA -3' |
|  | 3' - CCCGCCGGCGC -5' | 3' - TTTTTTTTTT -5' |

**Figure 1:**
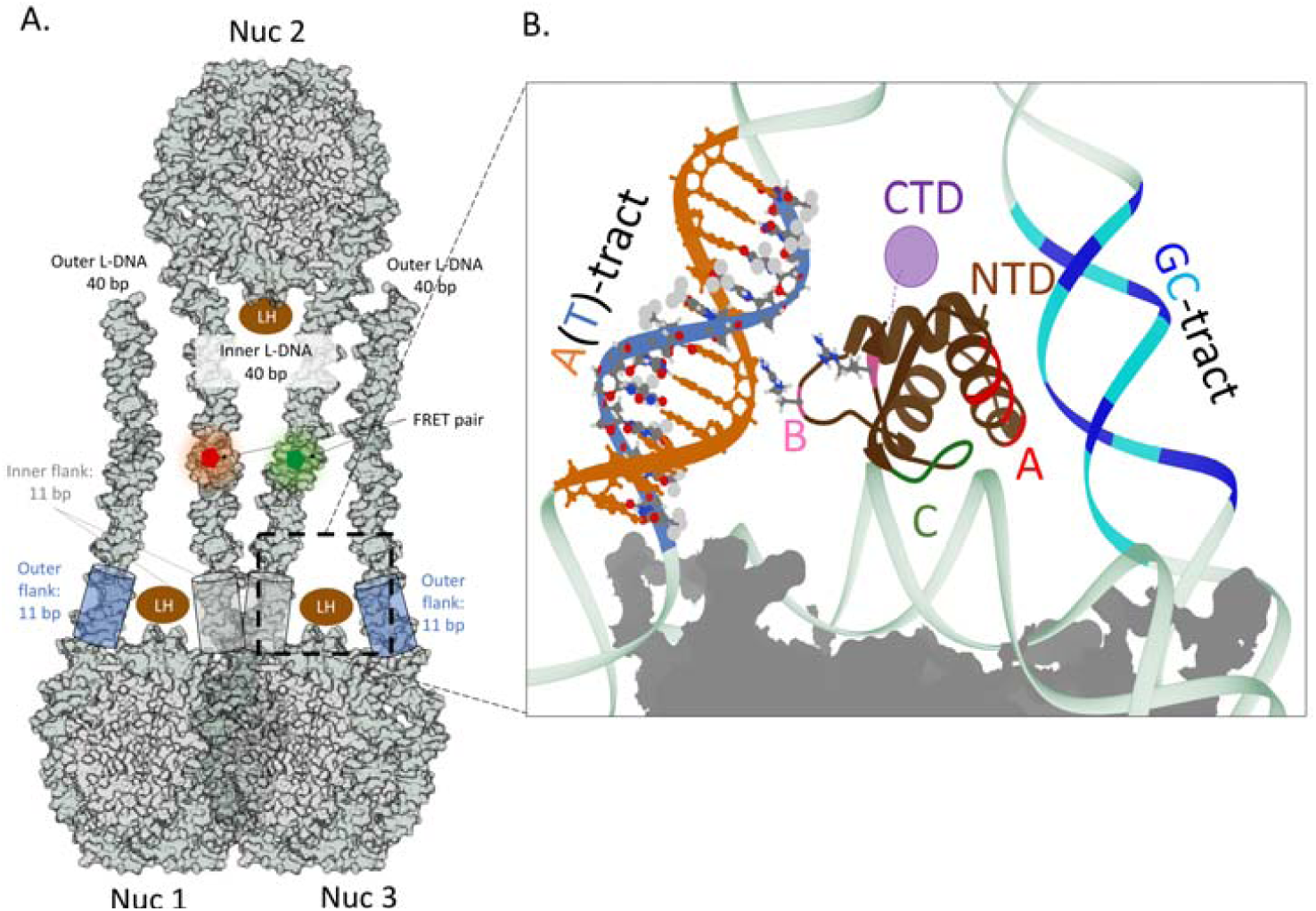
Structural features of the trinucleosome-LH systems studied. **A**. Schematic illustration of the trinucleosome systems studied. The three nucleosomes, Nuc 1, 2, and 3 are shown with the outer and inner L-DNAs labelled. The LHs are shown as brown ovals. The outer (blue) and inner (grey) flanks of the L-DNA arms are shown by cylinders. Fluorophores were attached to the methyl group of the thymine residue at the mid-point of each of the inner L-DNAs and are represented by green (Alexa 488 or donor) and red (Alexa 594 or acceptor) pentagons. **B**. Close-up of one of the nucleosomes showing the globular domain (gH) of the LH H1.0b (PDB ID 5NL0 (5)) positioned on-dyad with three DNA interacting zones, A (red), B (pink), and C (green) (27). Zone B can interact with and mediate recognition of A-tracts. The start of the C-terminal domain (CTD) is represented by a purple circle. The start of the N-terminal domain is labelled as NTD. The on-dyad gH is oriented towards the A-tract (dA in orange, dT in blue) minor groove on the L-DNA, with interactions mediated by two arginines in zone B (27). The gHs that bind to Nuc 1 and Nuc 3 in the trinucleosome constructs are expected to be similarly oriented to the A-tract flanking sequences.

## Materials and Methods

### Preparation of trinucleosomal DNA

For trinucleosome reconstitution, two 600 bp DNA sequences were constructed containing three Widom 601 sequences 40 bp apart and the respective inner and outer L-DNA flanking sequences. These constructs were built using a modified Golden-Gate assembly (29–31) *in vitro* that ensured that the three DNA fragments corresponding to the three nucleosomes were correctly oriented. In the Golden-Gate assembly protocol, the use of a Type IIs restriction enzyme (BsaI-HF-v2, New England Biolabs) that produces unique 4 bp sticky ends ensures that the three fragments are oriented in a unique way.

In detail, the *in vitro* Golden-Gate (29–31) assembly protocol consisted of: 1. PCR (polymerase chain reaction) of fragments 1, 2, 3 (fragment 1 and 3 purchased from Biolegio BV, Netherlands, and fragment 2 obtained from pGEM-3z/601 plasmid (28)) was performed separately to incorporate fluorophores into fragments 1 and 3, to incorporate a recognition site for BsaI-HF-v2 (New England Biolabs) in fragments 1, 2, and 3, and to amplify the fragments. The lengths of the PCR products were 242 bp (Fragments 1 and 3) and 172 bp (Fragment 2). Clean up of the PCR products was done using a standard cleanup protocol (NucleoSpin Gel and PCR cleanup, Macherey-Nagel GmbH). 2. Digestion reaction: The enzyme BsaI-HF-v2 (New England Biolabs) was used to digest the fragments at a concentration of 1.5 to 3 units / μg of DNA / 20 μl reaction volume. The time duration for the digestion reaction was between 18 and 36 hours. The reaction was carried out at 37°C. Digestion products and efficiency were checked by 8% polyacrylamide gel electrophoresis (Figure S1) and redigestion was performed if necessary. 3. Ligation reaction: The properly digested fragments were added in the following ratio of molecular weights: 1.00:0.65:1.25. 500 units of T4 ligase (New England Biolabs) / μg DNA / 300 μl volume were used. The duration of the ligation reaction was 24 hours at 15°C. 4. After ligation, the products were checked using 1% agarose gel electrophoresis. The 600 bp DNA was extracted from the gel using a standard NucleoSpin Gel and PCR cleanup (Macherey-Nagel GmbH). The fluorescence of the doubly labelled 600 bp band was visualized using a Typhoon 9400 scanner (GE Healthcare) and it was cut out.

### Reconstitution of trinucleosomes with and without LH

The reconstitution of the trinucleosomes was carried out similarly to the mononucleosome reconstitution described in De *et al*. (2021) (27), and involved a two-step dialysis method to bring down the concentration of NaCl from 2 M to about 25 mM. The molar ratio of octamers constructed from full-length, recombinant *Xenopus laevis* core histones (obtained from Planet Protein Colorado and purified following protocol referred earlier (32)) was 8 times that of the 600 bp DNA. The DNA and octamers were mixed in a TE buffer (10 mM Tris and 0.1 mM EDTA (ethylene diamine tetraacetic acid, pH 7.5) containing 2 M NaCl, by gently rocking at 4°C for 30 minutes. This reaction mix was transferred to a Slide-A-Lyzer MINI Dialysis unit (Thermo Fischer Scientific) with a cutoff of 7 kDa. With the help of a flotation device, this MINI dialysis unit was floated onto a second dialysis unit containing 15 ml of 2M NaCl TE solution. This second dialysis unit was floated onto a 1 litre TE buffer having 0.6 M NaCl. Dialysis was carried out for 7 hours at 4°C with slow stirring. This brought down the concentration of NaCl in the MINI dialysis unit from 2 M to about 0.6 M, enabling the reconstitution of the 600 bp DNA and the core histone octamers. After 7 hours, the LH (full-length *Xenopus laevis* H1.0b, wild-type recombinant, purified using the protocol described previously (27)) was added to the reaction mix in the MINI dialysis unit, at a LH:DNA ratio of 3.2. The two dialysis units were then refloated on 1 litre of TE buffer devoid of NaCl. The second step of dialysis was performed overnight, until the salt concentration of the reaction mix in the MINI dialysis unit was lowered to 25 mM NaCl.

Reconstitution was controlled by gel electrophoresis (Figure S1 ii) and by AFM imaging (Figure S1 iii). 1% agarose gel electrophoresis showed that the presence of LH leads to slower trinucleosome band migration (Figure S1 ii). AFM sample preparation and image acquisition were done as described before (33). Images were acquired in air using Nanoscope V (Digital Instruments) and Nanoscope software (v7.13). The AFM images verified the successful reconstitution of trinucleosomes (a representative trinucleosome is indicated by the white arrow for the A-near +LH system, Figure S1 iii).

### Single-pair FRET spectroscopy

The protocol for single-laser excitation FRET (single laser excitation with Cobolt Calypso 491 nm, Hübner Photonics, Germany) of doubly labelled samples and data acquisition have been described previously (34, 35). First, measurements were made for a donor-only (DNA labelled with the Alexa 488 donor dye, Figure S2A) sample to correct for potential crosstalk. Then, measurements were made for the doubly labelled reconstituted trinucleosomes at different salt concentrations (Figures S2B and S3 and Tables S4 and S5).

Single-laser excitation spFRET measurements were carried out for the freely diffusing trinucleosomes as described previously(27, 32, 34, 36–40). To study the stability and observe how the trinucleosomes dissociate with an increase in salt concentration, and how the presence of the LH affects such dissociation (41–43), experiments were performed in TE buffer at NaCl concentrations ranging from 25 to 700 mM NaCl. From spFRET measurements, we obtained the proximity ratio for each molecule, P, which is directly proportional to the FRET efficiency and computed by dividing the number of photons detected in the acceptor channel (N_A_) by the total number of photons detected in the acceptor and donor (N_D_) channels, i.e. P = N_A_/(N_A_+N_D_) (32, 37, 38). For each sample, proximity ratio histograms were fitted to multiple Gaussian peaks. The results of single-pair FRET measurements and fitting are shown in Figures S2 and S3, with the corresponding data (mean proximity ratio of major population peak(s), its relative population and its peak width at half maximum) in Tables S4 and S5, respectively. The proximity ratio histograms in Fig S2B and S3 (and fitted data in tables S4 and S5) show measurements taken at different NaCl concentrations: 25 mM (i), 75 mM (ii), 150 mM (iii), 250 mM (iv), 300 mM (v), 550 mM (vi) and 700 mM (vii). The fitting populations are shown with their peaks labelled as follows: Peak a: donor only population; peak(s) b: these peaks in the proximity ratio range of 0.1-0.2 denote partly dissociated products or free DNA; peak c: trinucleosome population (indicated by a black arrow); peak d (Figure S3): trinucleosome with LH (indicated by a black arrow). These peaks have a higher proximity ratio than for the trinucleosome without LH (peak c).

### Preparing the trinucleosome model

Since the proximity ratio profiles of A-near and A-far trinucleosomes with LH are highly similar (Figures 3, S6), only one trinucleosome was modeled with the mean proximity ratio of peak d at 25 mM NaCl (Figure 3, S6). Two previously published nucleosome structures (PDB id: 5NL0 (5), and 7K5X (6)) were first preprocessed by removing the LH and additional proteins. Using Chimera (44), their sequences were mutated to resemble that of Nuc 2 and its adjoining linker DNAs upto approximately 23 bp from the Widom 601 positioning sequence (also referred to as the core DNA sequence). The nucleosome structures were energy minimised with 100 steps of steepest descent (step size 0.02 Å) minimisation followed by 10 conjugate gradient steps (step size 0.02 Å), using the AMBER ff99bsc0 force field (45). Then these nucleosome structures were subjected to elastic network normal mode analysis (46) to generate a wide range of arm opening conformations. Using the software FRET Positioning and Screening (FPS) (47, 48), accessible volume simulated fluorophores were attached to the generated ensemble on the labelling positions, with dye parameters described previously (27, 47–49). Thus, the FPS software gave the computed proximity ratio between the fluorescent label binding sites (approximately 20 bp from the core DNA of Nuc 2). The conformer showing a proximity ratio of 0.8 was chosen to represent the middle or Nuc 2 nucleosome. Using Chimera, the two other nucleosomes, Nuc 1 and Nuc 3 (sequence changed to resemble Nuc 1 and Nuc 2 of A-far and A-near constructs and minimised following the same protocol as Nuc 2) were placed at the ends of the nucleosome 2 in such a way that the inner L-DNA is continuous and unbent. For simplicity and because of a lack of experimental data on their conformations, the outer L-DNA arms were truncated upto the flanks (denoted by blue cylinders in Figure 1) that either had an A-tract or GC-tract, depending upon the trinucleosome construct. By aligning, renumbering and renaming the three separate DNA pieces from the three nucleosomes on Chimera, one continuous DNA threading through the core octamers of Nuc 1, 2 and 3 was built. No further minimisations were done on the trinucleosome model. Minor clashes on the DNA, if present, were fixed manually using the move command in Chimera. After building the trinucleosome model, FPS was used to check once more that the computed FRET between the inner L-DNA arms matched the experimentally observed value of 0.8.

The gH domain was positioned on Nuc 2 in a canonical on-dyad fashion as observed in the structures: 4QLC (3), 5WCU (4), 5NL0 (5), and 7K5X (6). The gH domains were positioned on Nuc 1 and Nuc 3 according to the AG and GA models described in our earlier work (27), i.e., on-dyad, with Zone B proximal to the A-tract minor groove (as in Figure 1B ii). As a control, the gH was also positioned in an off-dyad position on Nuc 1 and Nuc 3 in the two trinucleosome structures to assess the possibility of direct gH-gH interactions (Figure S6i).

## Results and Discussion

### The linker histone H1.0b compacts trinucleosomes similarly irrespective of the placement of A-tract flanking sequences

We studied trinucleosome dissociation by measuring FRET at increasing NaCl concentrations to probe for eventual differences in their stability revealed by the dissociation pattern of the two types of trinucleosomes. Proximity ratio histograms calculated from the spFRET allows to distinguish subpopulations with different dye-dye distances. Such histograms obtained for the systems studied at seven different salt concentrations are shown in Figure 2 and Supplemental Figures S2-S3. The change in their pattern in both the LH-containing trinucleosomes (A-near and A-far) is similar at all salt concentrations, as shown in Figure 2A and B. In the absence of LH, the mean of the highest proximity ratio peak varies from 0.34 (low salt) to about 0.68 at 250 mM NaCl concentration (Figure 2B iii). With further increasing salt concentration, the mean proximity ratio of this peak falls to about 0.24 (700 mM), suggesting that dissociation of the trinucleosome occurs. In the presence of LH (Figure 2Bi and ii) and at NaCl concentrations from 25 mM to 300 mM, the highest proximity ratio peak is fit by two Gaussians (Supplemental figure S3) with comparable populations c and d, with d being the Gaussian with the higher proximity ratio). Population or peak d (black trace in Figure 2B i-ii) maintains a stable mean proximity ratio between 0.85 and 0.93 with a small peak width, indicating a highly compacted trinucleosome with restricted movement of the inner L-DNAs. While the LH is present in population c as well, as suggested by its high mean proximity ratio values at low salt concentration, this population could represent trinucleosomes having a somewhat relaxed compaction with the larger peak width indicating greater flexibility of inner L-DNA arms. Given that cryo-EM structures of 187 NRL nucleosome arrays have been observed to have LH depleted nucleosomes (51), the possibility that population or peak c contains less than three LHs per trinucleosome at low salt concentration cannot be discounted. Population d diminishes at 300 mM salt concentration, and at salt concentrations of 550 and 700 mM, trinucleosomes with and without LH all show only one population with a peak with a similar mean proximity ratio in the presence and absence of LH. This suggests that the LH starts to dissociate at salt concentrations of 300 mM and completely dissociates at salt concentrations of 550 mM and 700 mM.

**Figure 2:**
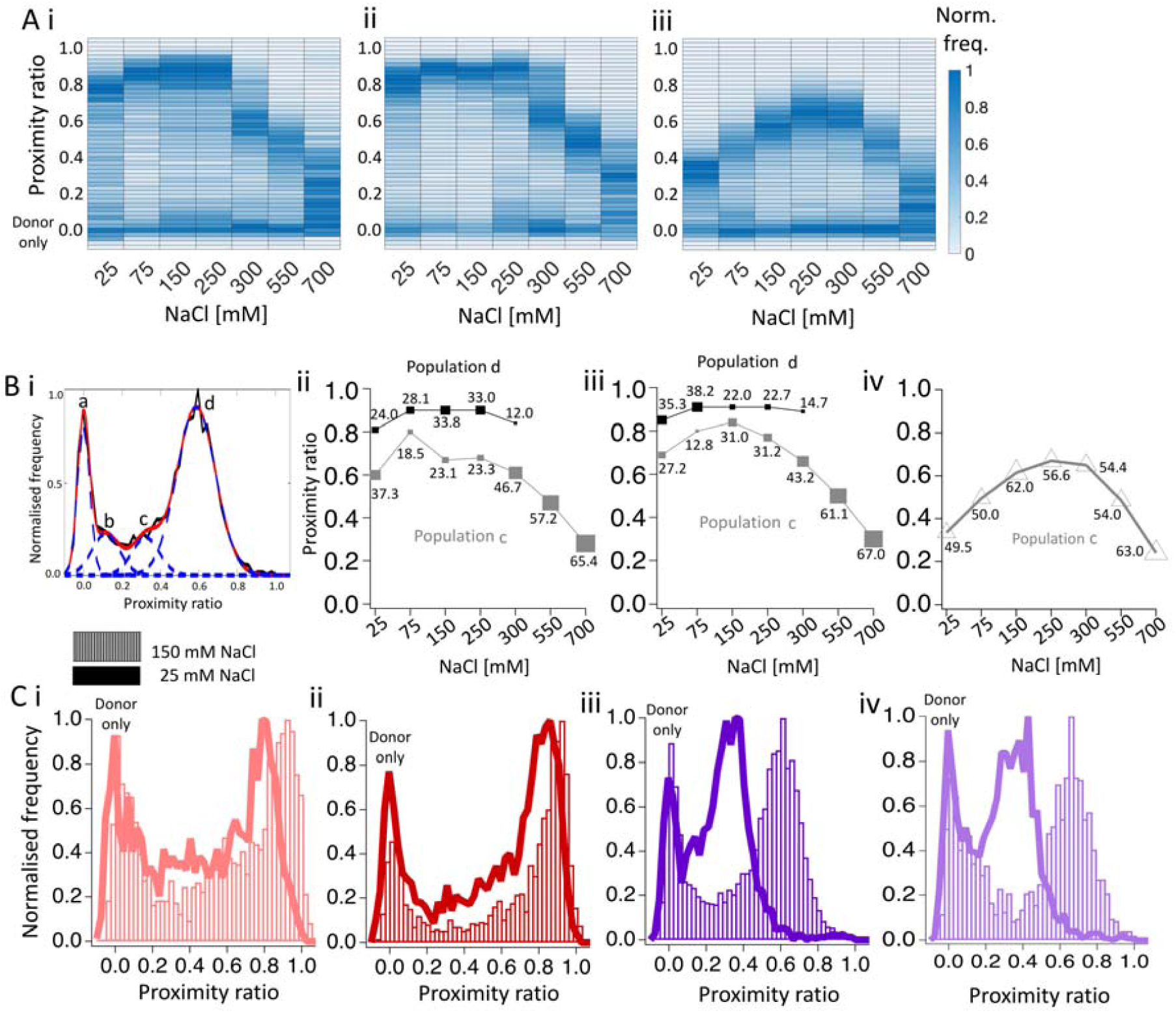
spFRET measurements of the compaction of the trinucleosomes, A-far and A-near, with and without LH. **A**. Heat map representations of proximity ratio histograms measured by spFRET under seven NaCl concentrations ranging from 25 to 700 mM for **(i)** A-far with LH, **(ii)** A-near with LH, and **(iii)** A-near without LH. The colour intensity represents the normalised frequency of the proximity ratio value. **B.(i)** A representative proximity ratio histogram deconvoluted into 4 populations: ‘a’, ‘b’, ‘c’ and ‘d’. (ii-iv) Plots of mean proximity ratio of populations c and d of the high proximity ratio major population peaks (see Supplemental Figure S2 and S3) against salt concentration. The marker size and label show the % population of the peak. **(ii)** A-far with LH, **(iii)** A-near with LH, **(iv)** A-near without LH. The proximity ratio histograms were similar for A-far without LH to those for A-near without LH (see Supplemental Figure S2). **C**. Proximity ratio (P) histograms measured at 25 mM (solid line) and 150 mM (bars) NaCl concentration for **(i)** A-far with LH, **(ii)** A-near with LH, **(iii)** A-near without LH, and **(iv)** A-far without LH. (Histograms measured for other salt concentrations are given in see Supplemental Figure S2 and S3).

For the trinucleosomes with LH (Figure 2.A and Bi and ii), the mean proximity ratios of the selected peaks are higher than those without LH (Figure 2.A and B iii), at salt concentrations from 25 mM to 250 mM. This difference confirms that the LH compacts the trinucleosomes, bringing the inner L-DNA arms closer to each other. Shielding of the negative charges on the DNA backbone by the positively charged sodium ions also brings the two inner L-DNA arms closer to each other in trinucleosomes lacking LH (Figure 2.Biii). This increase in compaction as a result of charge shielding by the salt is more pronounced in trinucleosomes lacking LH (Figure 2C iii and iv) than trinucleosomes associated with LHs (Figure 2C i and ii).

Our results indicate that the LH compacts the A-near and A-far trinucleosomes similarly at 25 mM salt. The proximity ratio histograms for the other salt concentrations show that this similarity between A-near and A-far trinucleosomes holds over the range of salt concentrations studied, their stability is unchanged.

Next, we focused on population d, the Gaussian peaks with the higher proximity ratios and lower widths, to investigate what three-dimensional structure it corresponds to and why, despite the different positions of the A-tracts, LH binding does not appear to affect the relative extent of compaction of the A-far and A-near trinucleosomes. The mean proximity ratio of population d only varies from 0.8 to 0.9, indicating little variation in the interfluorophore distance over the population (51). Thus, for each of the two DNA sequences studied, we built a single representative trinucleosome model that has a computed FRET efficiency between the inner L-DNA arm fluorophores of 0.8.

### The modeled structures of LH-bound trinucleosomes with a high FRET efficiency correspond to a 2-start helical arrangement

As seen in Figure 3i, the trinucleosome model (the DNA sequence of the model shown corresponds to A-near) corresponding to population d at low salt shows a standard 2-start helical arrangement (52) as predicted and observed for 187 NRL nucleosomes (53–55). The stacking between nuc 1 and nuc 3 is good as confirmed by aligning the modelled structures (for A-near and A-far) onto the experimentally determined structure of a stacked LH-bound 187 NRL tetranucleosome (PDB ID: 7PF2 (50)) (Figure S6ii), and a LH-bound hexanucleosome (PDB ID 6HKT (54) Figure S6iii).

**Figure 3:**
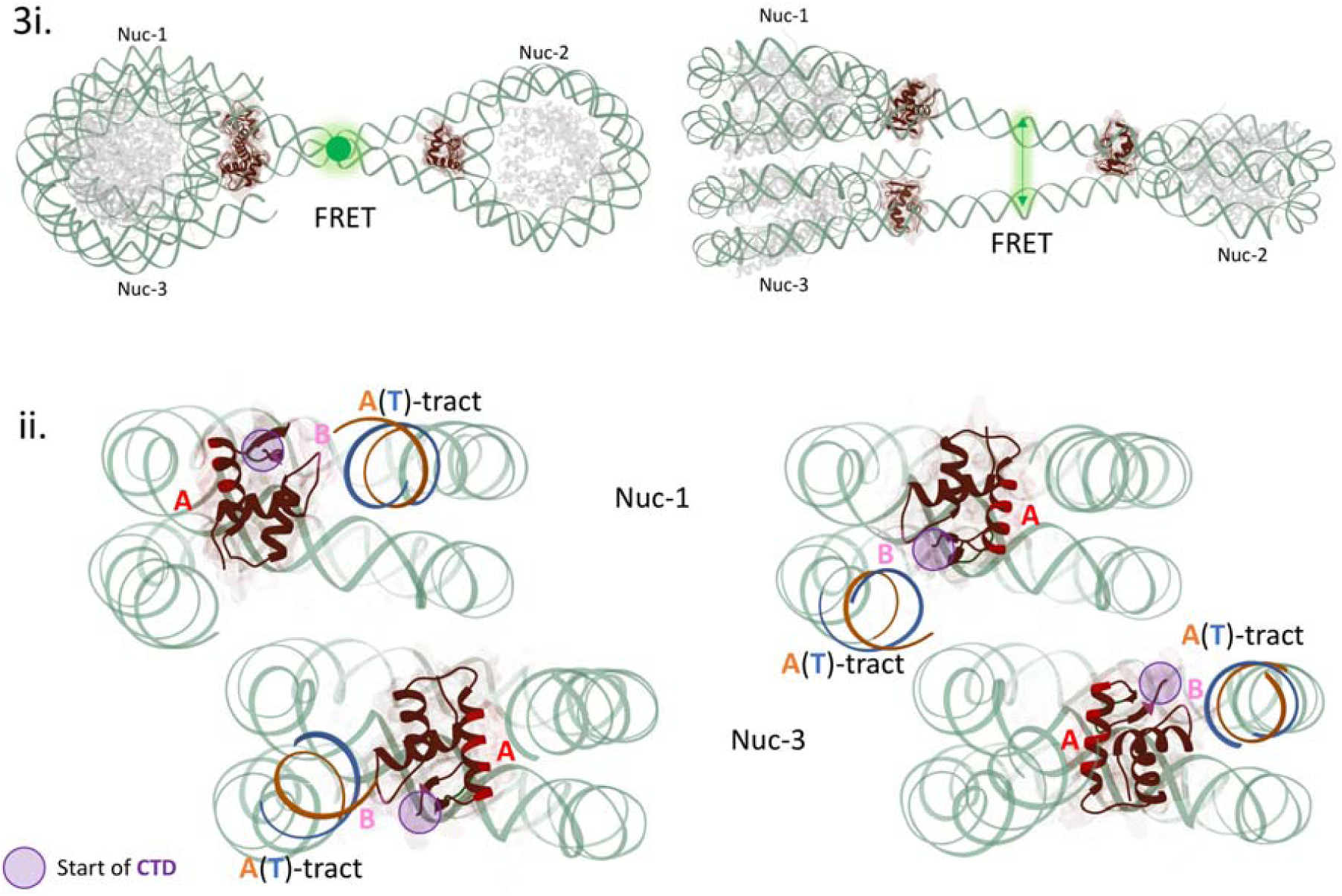
Model of a representative structure of population d of the A-near and A-far LH-bound trinucleosomes. (i) Two perpendicular views of a model with a computed FRET efficiency between the labels on the inner L-DNA arms of 0.8. The outer L-DNAs were truncated for clarity. The green arrows represent energy transfer between the fluorophore labels (which are not shown). Nuc 1 and Nuc 3 stack. (ii) Stacking of LH-bound Nuc 1 and Nuc 3 showing the different relative positioning of the on-dyad gHs. Left: A-near and Right: A-far. Each LH is shown with its gH positioned on-dyad and oriented towards the A-tracts (27). The LH gH is colour-coded as in Figure 1. The core histone octamers are shown in grey, the DNA is pale green. The A-tracts are coloured with adenines in orange and thymines in blue. The start of the LH CTD is denoted by a purple circle.

To understand why, despite having A-tracts at different positions (either on the inner or on the outer L-DNA arms of Nuc 1 and Nuc 3), the two trinucleosomes were compacted similarly by the LH, we focused on the interactions of the gHs on Nuc 1 and Nuc 3.

### The orientations of the on-dyad linker histones on stacking nucleosomes do not influence trinucleosome compaction

Although we did not directly observe the location of the full-length LH in our experiments, we assume that the gHs (of LH of subtype H1.0b) associate in an on-dyad position (5, 6, 56) in an orientation like the one that we observed for A-tract-containing mononucleosomes (27), shown in Figure 1Bii. The models for A-near and A-far with the LHs positioned in this way are shown in Figure 3ii. In A-near (Figure 3ii left), the A-tracts are on the inner L-DNA and in A-far, they are on the outer L-DNA (Figure 3ii right). For the gH to affect compaction of a nucleosomal array, it would be expected to self-associate (8) or form salt-bridges. In our models, the on-dyad gH domains on Nuc 1 and Nuc 3 are not close enough to each other to self-associate. These models suggest that the orientation of the gH domains of the LH H1.0b does not play a role in the compaction of the trinucleosomes.

We next considered the role of the LH CTDs. The purple circles in Figure 3ii represent the C-terminus of the gH domain, corresponding to the start of the 100-residue long intrinsically disordered CTD tail. If the CTDs associate with the L-DNA arm to which the C-terminus of the gH (purple circle) is closest, then the CTD would associate with the inner L-DNA arm of A-near and with the outer L-DNA arm of A-far. In this case, we would expect that the highly positively charged CTD would bring the inner L-DNAs closer together in A-near than A-far, resulting in a higher FRET efficiency between the fluorophores.

In our previous work on mononucleosomes, we have observed that position 101 (6 residues into the CTD from the gH) is nearly equidistant from both L-DNA arms (27). Zhou *et al*. (2021) (6) observed that the CTD of LH H1.0b (PDB 7K5X) associates with both L-DNA arms. But coarse-grained simulations have shown that the CTD can associate with one L-DNA at low salt (5 and 15 mM), and with both L-DNA arms at high salt (80 and 150 mM) concentrations (9). If this is the case, then we would expect that, at low salt concentrations, A-near would be more compact compared to A-far. In Figure 2B, the mean proximity ratio peak of Population c of A-near with LH is slightly higher than A-far with LH at salt concentrations of 25, 150 and 250 mM (difference, ΔP_mean_, of about 0.1, Supplemental tables S4 and S5), whereas the more compact Population d shows similar mean proximity ratios (ΔP_mean_ ∼ 0.05) for both A-near and A-far. This difference in proximity ratio would only correspond to a small difference (about 0.1 to 0.2 nm) in interfluorophore distance. Thus, these data provide little or no support for a strong association of the CTD with the inner L-DNA arms of A-near.

Overall, considering both populations c and d, the mean proximity ratios of the two trinucleosome sequences remain similar as the salt concentration is raised, suggesting that the LH CTDs associate with both inner and outer L-DNA arms of Nuc 1 and Nuc 3 and, therefore, contribute similarly to the compaction of the two trinucleosomes, despite the different locations of the A-tracts.

Although we observe here that the gH domains of on-dyad positioned LH are not sufficiently close to dimerise or form salt bridges, we considered whether an off-dyad positioning (to A-tract containing arm) gH would be close enough. Figure S6i shows a model where the off-dyad gH are close enough to dimerise or form salt bridges in the A-near model, and far apart in A-far. We suggest that, had the gH associated off-dyad, A-near could have been more compact than A-far. Since the compaction of A-near and A-far are similar, the gH likely associates on-dyad.

## Conclusions

We find that the different locations of the A-tracts in the DNA sequences of the two trinucleosomes studied do not affect LH-mediated trinucleosome compaction (Figure 2), despite the expected A-tract-mediated orientation of the on-dyad LH gH domains (27) (Figure 1B ii). Modeling constrained by the experimental observations (Figure 3) indicates that this is because the gH domains of the stacked nucleosomes in the trinucleosomes are too far away from each other to associate. Furthermore, analysis of the spFRET data at different salt concentrations indicates that the LH CTDs contribute to trinucleosome compaction but that their interaction with the L-DNA arms of the nucleosomes is rather insensitive to the locations of the inserted A-tracts. These results thus point to a robustness of chromatin structure to heterogeneity at the level of linker DNA sequence and individual nucleosomes. However, further study would be necessary to assess the generality of our observations for other DNA and linker histone sequences.

Here, we studied the LH H1.0b isoform, whose gH has been observed to associate in an energetically favourable on-dyad fashion (5, 6, 13, 56). A different LH isoform with an off-dyad gH binding mode or an altered affinity, due to differences in sequence or post-translational modification in the gH or CTD, may show a different sensitivity to L-DNA sequence and A-tract insertions, as regards its ability to compact chromatin (57). Likewise, less strongly nucleosome positioning sequences than the Widom 601 sequences studied here may result in altered sensitivity of LH-mediated compaction to A-tract insertions in the L-DNA. Furthermore, we studied two sequences with differently positioned A-tracts but clearly, further variations on DNA position and sequence could be studied, and additional positions for fluorophore labelling could be used to provide more structural constraints. Nevertheless, this study shows how the preference of a LH for A-tracts can be exploited to probe the determinants of chromatin compaction, as well as to understand to what extent LHs – as ubiquitous proteins – regulate chromatin structure, and thus transcription.

## Abbreviations

spFRET: single pair Förster/Fluorescence Resonance Energy Transfer
L- DNA: linker DNA
LH: Linker histone
gH: globular (head) domain of the LH
CTD: C-terminal domain of the LH
AFM: Atomic Force Microscopy

## Supporting Material

consists of details of the methods (Fig S1), proximity ratio histograms (Fig S2, S3, Tab. S4, S5) and additional modelling data (Fig S6). The spFRET data are available at https://doi.org/10.5281/zenodo.5090767

## Acknowledgements

The authors are very grateful to Prof. Jeffrey Hayes and Dr. Amber Cutter for providing us with the plasmid for wild-type linker histone. The authors are grateful to Nathalie Schwarz for purifying the linker histone. This work was supported by the Helmholtz International Graduate School for Cancer Research (DKFZ) and the Klaus Tschira Foundation.

## Author contributions

MD, KT and RCW conceived the research. MD, GM, KT designed the DNA constructs. MD and GM prepared and purified the DNA. MW acquired the AFM images. MD performed the spFRET measurements with suggestions from KT, and the modeling with suggestions from RCW, and wrote the paper with suggestions from all the authors. All the authors contributed to the finalisation of the manuscript.

