## Supporting Material for "Robustness of trinucleosome compaction to A-tract mediated linker histone orientation"

### Materials and Methods

#### Preparation of trinucleosomal DNA

**A. Primer sequence**

Unlabelled and labelled primers were purchased from Sigma-Aldrich (USA) and IBA Lifesciences (Germany), respectively. All primers are listed from 5′ to 3′ ends. Overhangs allowing BsaI-HF-v2 to properly dock onto the recognition site are shown in italic. Recognition sites for BsaI-HF-v2 are shown in bold. The cutting site is denoted by a '|'. The 4 bp unique sticky ends left after digestion (denoted as NNNN and FFFF in Figure S1) that enable ligation in a single orientation are underlined. The thymines labelled with fluorophores are shown in green (Alexa 488) and red (Alexa 594).

Nuc 1 Forward unlabelled: ATCCGACTGGCACCGGCAATGTCGCTG

Nuc 1 Reverse-Alexa 488: *CCATC***GGTCTC**ACTGT|GAGTTCATCCCTTATGTGA**T**GGACCC

Nuc 2 Forward unlabelled: *ATCGA***GGTCTC**T|ACAGGATGTATATATCTGACACGTGCC

Nuc 2 Reverse unlabelled: *CCATC***GGTCTC**ACCTG|GAGAATCCCGGTGCC

Nuc 3 Forward-Alexa 594: *GTCAT***GGTCTC**T|CAGGATCCGACTGGCACCGGCAA**T**GTCGC

Nuc 3 Reverse unlabelled: GAGTTCATCCCTTATGTGATGGACCCT

**B. Trinucleosome DNA sequence**

Two 600 bp trinucleosome sequences were generated named A-near and A-far according to the positioning of the A-tracts. In A-near, the A-tracts were on the inner linker-DNA arms of nucleosomes 1 and 3. In A-far, the A-tracts were on the outer-DNA arms of nucleosomes 1 and 3. The inner and outer flanking sequences are underlined. The bases denoted in mauve italics denote the Widom 601 core nucleosome positioning sequence. The green and red coloured bases indicate the positions of the fluorophore labels: green adenine denoted in bold: Alexa 488 on the thymine residue of the complementary strand; red: thymine labelled with Alexa 594.

**A-near:**

ATCCGACTGGCACCGGCAATGTCGCTGTTTACGCGGCCGCCC*TGATGTATATATCTGACACGTGCCTGGAGACTAGGGAGTAATCCCCTTGGCGGTTAAAACGCGGGGGACAGCGCGTACGTGCGTTTAAGCGGTGCTAGAGCTGTCTACGACCAATTGAGCGGCCTCGGCACCGGGATTCTCCC*TTTTTTTTTTTGGTAGGGTCC**A**TCACATAAGGGATGAACTCACA*GGATGTATATATCTGACACGTGCCTGGAGACTAGGGAGTAATCCCCTTGGCGGTTAAAACGCGGGGGACAGCGCGTACGTGCGTTTAAGCGGTGCTAGAGCTGTCT*

*ACGACCAATTGAGCGGCCTCGGCACCGGGATTCTCC*AGGATCCGACTGGCACCGGCAA**T**GTCGCTGTTCCAAAAAAAAAAA*GGATGTATATATCTGACACGTGCCTGGAGACTAGGGAGTAATCCCCTTGGCGGTTAAAACGCGGGGGACAGCGCGTACGTGCGTTTAAGCGGTGCTAGAGCTGTCTACGACCAATTGAGCGGCCTCGGCACCGGGATTCTCCA*GGGCGGCCGCGTATAGGGTCCATCACATAAGGGATGAACTC

**A-far:**

ATCCGACTGGCACCGGCAATGTCGCTGTTCCAAAAAAAAAAA*GGATGTATATATCTGACACGTGCCTGGAGACTAGGGAGTAATCCCCTTGGCGGTTAAAACGCGGGGGACAGCGCGTACGTGCGTTTAAGCGGTGCTAGAGCTGTCTACGACCAATTGAGCGGCCTCGGCACCGGGATTCTCCA*GGGCGGCCGCGTATAGGGTCC**A**TCACATAAGGGATGAACTCACA*GGATGTATATATCTGACACGTGCCTGGAGACTAGGGAGTAATCCCCTTGGCGGTTAAAACGCGGGGGACAGCGCGTACGTGCGTTTAAGCGGTGCTAGAGCTGTCTACGACCAATTGAGCGGCCTCGGCACCGGGATTCTCCA*GGATCCGACTGGCACCGGCAA**T**GTCGCTGTTTACGCGGCCGCCC*TGATGTATATATCTGACACGTGCCTGGAGACTAGGGAGTAATCCCCTTGGCGGTTAAAACGCGGGGGACAGCGCGTACGTGCGTTTAAGCGGTGCTAGAGCTGTCTACGACCAATTGAGCGGCCTCGGCACCGGGATTCTCCC*TTTTTTTTTTTGGTAGGGTCCATCACATAAGGGATGAACTC

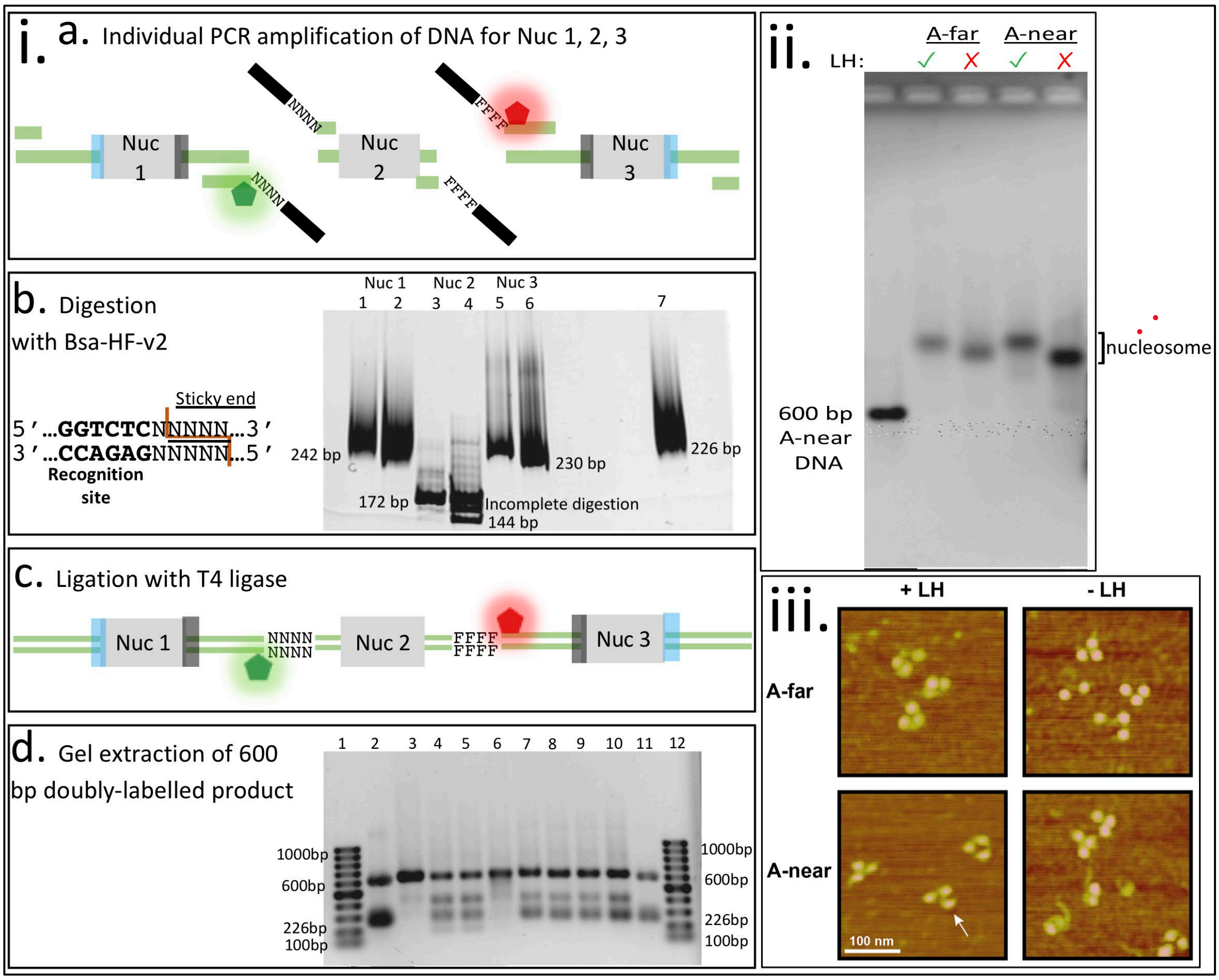

***Figure S1:*** *(i). Trinucleosome DNA preparation and reconstitution. i.a-d Steps of the modified in vitro Golden Gate Assembly protocol used to synthesize 600 bp DNA for the two trinucleosomes. Step a: PCR amplification of the DNA of the three nucleosomes (Nuc). Each Widom 601 [1] positioning sequence is denoted in grey with the flank sequences on Nuc 1 and Nuc 3 in blue (outer), or dark grey (inner). Green and red pentagons denote Alexa 488 and 594, respectively. 'NNNN' and 'FFFF' denote unique 4 bp sticky ends generated by BsaI-HF-v2 digestion. Step b: The DNA recognition site and sticky ends for BsaI-HF-v2 digestion and 8% polyacrylamide gels (120 V, 17 V/cm) run for 1.5h which show BsaI-HF-v2 digested (Lanes 2,6), undigested (Lanes 1,3,5), incompletely digested (Lane 4) bands. Step c: Ligation of the DNA to form the trinucleosome sequences was performed with T4 ligase. Step d: A comparison of extracted 600 bp and reaction mix seen in 1% agarose gel run at 116 V for 1 h. Lanes 1 and 12: 1 kbp ladders, 2 and 11: mix of 600 bp and 226 bp DNA, 3: A-far DNA 600 bp, 4 and 5: A-far preparation mix showing fully ligated 600 bp DNA, and incompletely ligated or unligated fragments of sizes 150, 200, 400 bp, 6: 600 bp A-near DNA, 7-10: A-near DNA preparation mix showing ligated and unligated products.* (ii and iii). *Trinucleosomes after reconstitution. (ii). 1% agarose gel (stained with ethidium bromide) showing reconstituted trinucleosomes with (green tick) and without (red cross) LH. Upon addition of the LH, the trinucleosome band migrates slower. The 600 bp A-near free DNA is shown in the 1st lane and migrates fastest. The reconstituted trinucleosome samples do not have any free DNA. (iii). Representative sections of atomic force microscopy (AFM) images of reconstituted trinucleosomes (highlighted with white arrow) for A-far and A-near with (+LH) and without LH (-LH). Although mono- and dinucleosomes were also observed, the major population was comprised of trinucleosomes.*

### Single-pair FRET spectroscopy

2A.

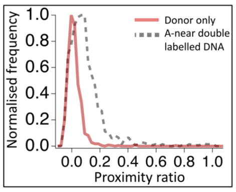


2B.

**Without linker histone**

A-near

A-far

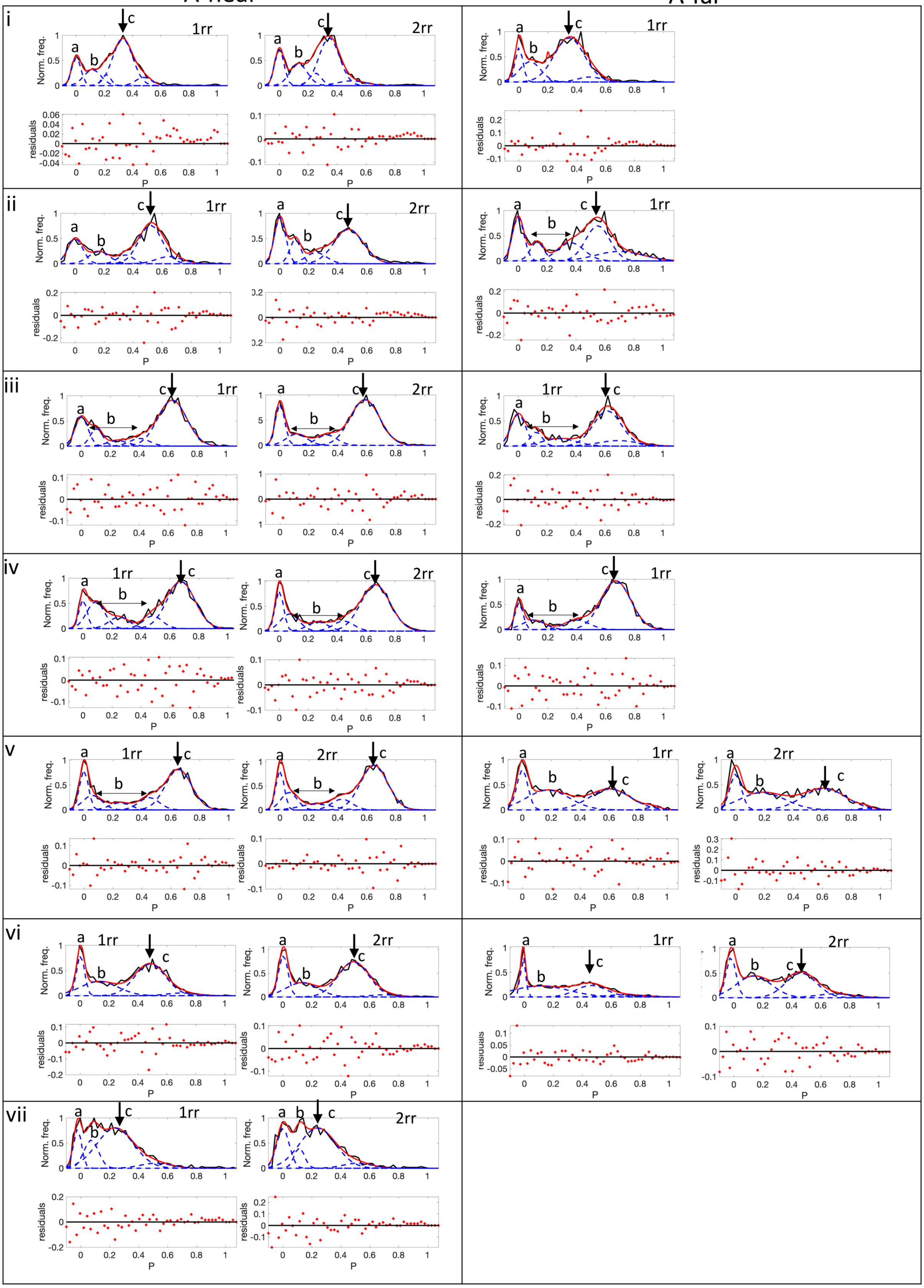

***Figure S2:*** *Single-pair FRET measurements. A. A donor-only (DO) sample (red) containing Alexa 488 dye was measured prior to experiments, to correct for cross-talk. Dashed line: The double-labelled A-near DNA shows a low proximity ratio peak at 0.1. B. Proximity ratio histograms of A-near and A-far trinucleosomes without LH at NaCl concentrations from 25 mM to 700 mM (for reconstitution replicates 1rr and 2rr), with plots of residuals from peak fitting below. Black: measured data, blue dashed line: fitting populations, red trace: fitted data. See Table S4 for fitted values.*

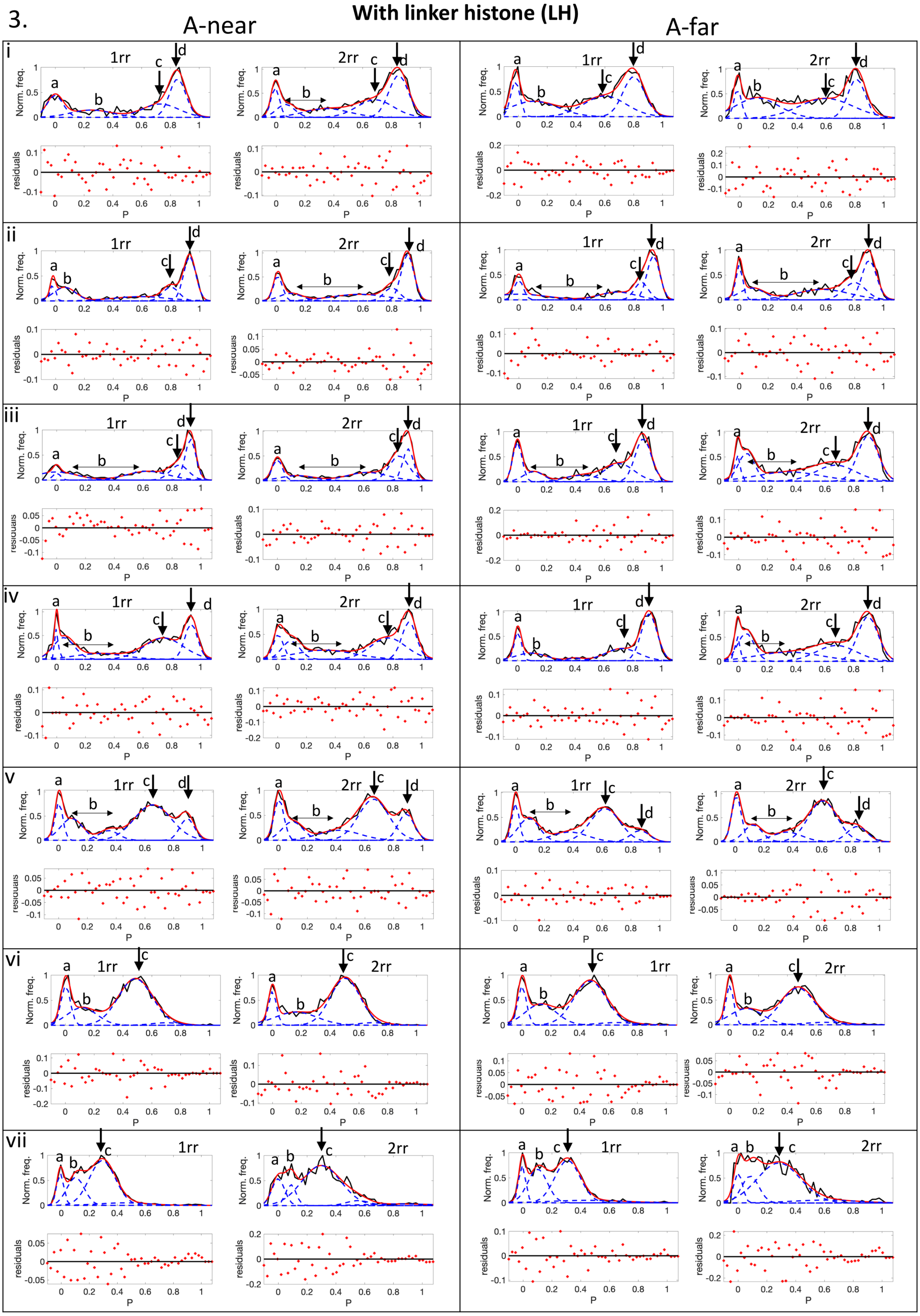


***Figure S3:*** *Proximity ratio histograms from single-pair FRET measurements of A-near and A-far trinucleosomes with LH, with plots showing residuals from peak fitting below. Black: measured data; blue dashed lines: fitting populations; red traces: fitted data. See Table S5 for fitted values.*

**Table S4:** Values from proximity ratio plot fitting shown in Figure S2B: mean proximity ratio (P) of peaks c, relative population, and full-width at half maximum (FWHM).

<table>
<tr><th rowspan="2">NaCl (mM)</th><th rowspan="2">Reconstit-ution Replicate (rr)</th><th colspan="3">A-near without linker histone</th><th colspan="3">A-far without linker histone</th></tr>
<tr><th>$P_{mean}$ (c)</th><th>% population</th><th>FWHM</th><th>$P_{mean}$ (c)</th><th>% population</th><th>FWHM</th></tr>
<tr><td rowspan="2">25</td><td>1rr</td><td>0.33</td><td>55.2</td><td>0.24</td><td>0.35</td><td>61.6</td><td>0.35</td></tr>
<tr><td>2rr</td><td>0.34</td><td>49.5</td><td>0.24</td><td colspan="3">No data</td></tr>
<tr><td rowspan="2">75</td><td>1rr</td><td>0.52</td><td>48.5</td><td>0.27</td><td>0.55</td><td>33.2</td><td>0.27</td></tr>
<tr><td>2rr</td><td>0.47</td><td>50.0</td><td>0.34</td><td colspan="3">No data</td></tr>
<tr><td rowspan="2">150</td><td>1rr</td><td>0.63</td><td>61.0</td><td>0.33</td><td>0.62</td><td>47.2</td><td>0.32</td></tr>
<tr><td>2rr</td><td>0.60</td><td>62.0</td><td>0.34</td><td colspan="3">No data</td></tr>
<tr><td rowspan="2">250</td><td>1rr</td><td>0.68</td><td>62.5</td><td>0.32</td><td>0.67</td><td>68.5</td><td>0.34</td></tr>
<tr><td>2rr</td><td>0.66</td><td>56.6</td><td>0.34</td><td colspan="3">No data</td></tr>
<tr><td rowspan="2">300</td><td>1rr</td><td>0.65</td><td>51.0</td><td>0.30</td><td>0.61</td><td>39.2</td><td>0.47</td></tr>
<tr><td>2rr</td><td>0.65</td><td>54.4</td><td>0.31</td><td>0.61</td><td>43.1</td><td>0.50</td></tr>
<tr><td rowspan="2">550</td><td>1rr</td><td>0.48</td><td>51.0</td><td>0.35</td><td>0.46</td><td>28.2</td><td>0.35</td></tr>
<tr><td>2rr</td><td>0.49</td><td>54.0</td><td>0.35</td><td>0.46</td><td>36.6</td><td>0.35</td></tr>
<tr><td rowspan="2">700</td><td>1rr</td><td>0.24</td><td>62.8</td><td>0.42</td><td></td><td></td><td></td></tr>
<tr><td>2rr</td><td>0.24</td><td>63.0</td><td>0.41</td><td></td><td></td><td></td></tr>
</table>

**Table S5:** Values from proximity ratio plot fitting shown in Figure S3: mean proximity ratio (P) of peaks c and d, relative population, and full-width at half maximum (FWHM).

| NaCl (mM) | Reconstit-ution Replicate (rr) | A-near with linker histone | | | A-far with linker histone | | |
|---|---|---|---|---|---|---|---|
| | | $P_{mean}$ (c,d) | % popul-ation (c,d) | FWHM (c,d) | $P_{mean}$ (c,d) | % popul-Ation (c,d) | FWHM (c,d) |
| 25 | 1rr | 0.73, 0.85 | 27.5, 35.6 | 0.40, 0.20 | 0.60, 0.80 | 31.3, 31.0 | 0.48, 0.25 |
| | 2rr | 0.69, 0.85 | 27.2, 35.3 | 0.40, 0.20 | 0.60, 0.81 | 37.3, 24.0 | 0.63, 0.20 |
| 75 | 1rr | 0.80, 0.93 | 15.8, 39.4 | 0.20, 0.14 | 0.85, 0.93 | 15.2, 35.0 | 0.16, 0.14 |
| | 2rr | 0.80, 0.91 | 12.8, 38.2 | 0.17, 0.15 | 0.80, 0.90 | 18.5, 28.1 | 0.25, 0.17 |
| 150 | 1rr | 0.85, 0.93 | 18.8, 30.0 | 0.21, 0.12 | 0.70, 0.87 | 27.0, 34.4 | 0.35, 0.20 |
| | 2rr | 0.84, 0.91 | 31.0, 22.0 | 0.22, 0.12 | 0.67, 0.90 | 23.1, 33.8 | 0.45, 0.23 |
| 250 | 1rr | 0.74, 0.93 | 39.2, 21.1 | 0.45, 0.14 | 0.72, 0.91 | 24.6, 45.8 | 0.38, 0.20 |
| | 2rr | 0.77, 0.91 | 31.2, 22.7 | 0.40, 0.17 | 0.68, 0.90 | 23.3, 33.0 | 0.46, 0.23 |
| 300 | 1rr | 0.65, 0.89 | 50.5, 12.1 | 0.43, 0.17 | 0.62, 0.86 | 42.1, 9.2 | 0.34, 0.22 |
| | 2rr | 0.66, 0.89 | 43.2, 14.7 | 0.35, 0.20 | 0.61, 0.84 | 46.7, 12.0 | 0.33, 0.23 |
| 550 | 1rr | 0.49 | 62.2 | 0.40 | 0.47 | 57.0 | 0.38 |
| | 2rr | 0.50 | 61.1 | 0.34 | 0.47 | 57.2 | 0.39 |
| 700 | 1rr | 0.30 | 58.0 | 0.31 | 0.31 | 49.2 | 0.30 |
| | 2rr | 0.30 | 67.0 | 0.45 | 0.28 | 65.4 | 0.50 |

### Preparing the trinucleosome model

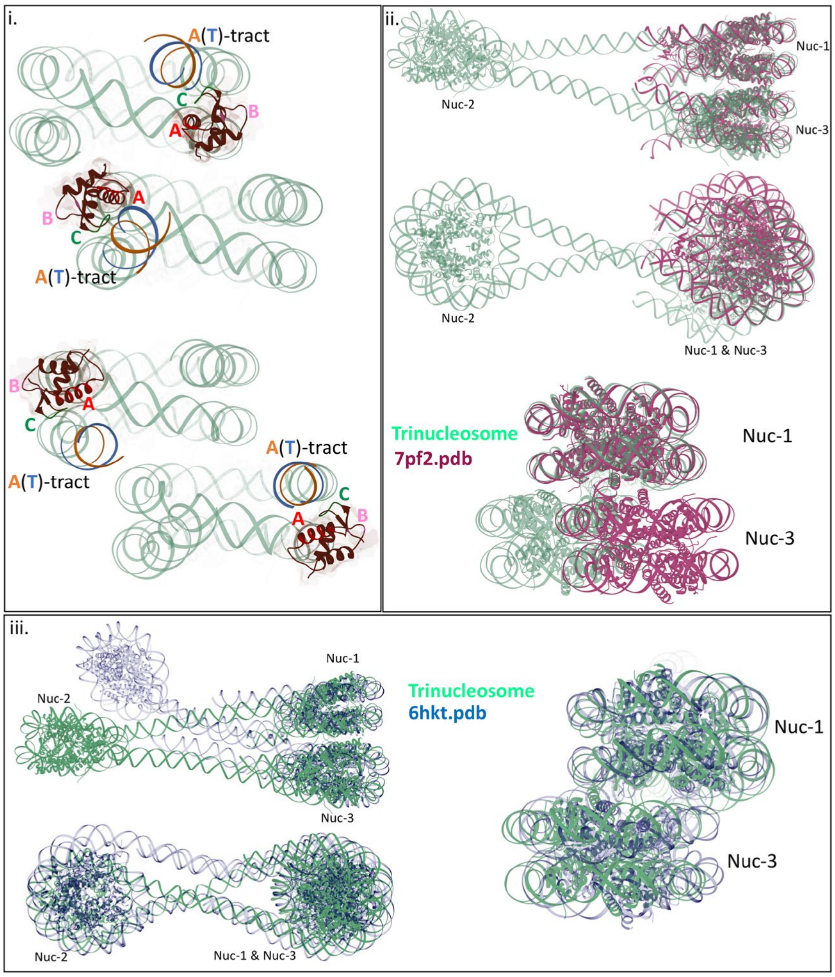


***Figure S6: Modeling of the trinucleosome-LH structures*** *(i) Trinucleosome models with the gH LH domains binding to nucleosomes Nuc 1 and 3 positioned off-dyad for the A-near (top) and A-far (bottom) structures. In the case of A-near, the LH gH domains positioned off-dyad might approach close enough to form salt bridges (not shown in the picture), see text for details (ii) Nuc 1 and Nuc 3 of the trinucleosome model (green trace) stack in a favorable arrangement as is shown by their overlay onto the cryo-EM structure of the stacked 187 NRL tetranucleosome containing human LH H1.4 (PDB id 7PF2[2]) shown in magenta. (iii) Nuc 1 and Nuc 3 of the general trinucleosome model (green trace) stack in a favorable arrangement as is shown by their overlay onto the crystal structure of the stacked H1-bound hexanucleosome (PDB id 6HKT [3]) shown in blue. For simplicity, only three nucleosomes of the hexanucleosomal array are shown.*
